# Rapid homeostatic modulation of transsynaptic nanocolumn rings

**DOI:** 10.1101/2021.06.15.448550

**Authors:** Paola Muttathukunnel, Patrick Frei, Sarah Perry, Dion Dickman, Martin Müller

## Abstract

Robust neural information transfer relies on a delicate molecular nano-architecture of chemical synapses. Neurotransmitter release is controlled by a specific arrangement of proteins within presynaptic active zones. How the specific presynaptic molecular architecture relates to postsynaptic organization, and how synaptic nano-architecture is transsynaptically regulated to achieve stable synaptic transmission remains enigmatic. Using time-gated stimulated emission depletion (gSTED) microscopy at the *Drosophila* neuromuscular junction, we here find that presynaptic nano-rings formed by the active-zone scaffold Bruchpilot (Brp) precisely align with postsynaptic glutamate receptor (GluR) rings. Individual rings harbor ~5 transsynaptically-aligned Brp-GluR ‘nanocolumns’. Intriguingly, acute GluR impairment rapidly triggers the formation of new transsynaptic nanocolumns on the minute time scale during homeostatic plasticity. We reveal distinct phases of structural transsynaptic homeostatic plasticity, with postsynaptic reorganization preceding presynaptic modulation. Finally, the auxiliary GluR subunit Neto-β promotes structural and functional homeostatic plasticity. Thus, transsynaptic nanocolumns arrange in stereotypic rings that are rapidly modulated during homeostatic plasticity to stabilize synaptic efficacy.

**One Sentence Summary:** Rapid changes of transsynaptic nanocolumn rings during homeostatic plasticity.

## Introduction

Even very subtle changes in the molecular architecture of chemical synapses may profoundly affect neural information processing and animal behavior^2,3^. Yet, neural systems are stable for a lifetime, implying robust maintenance of synaptic signaling. Synaptic transmission is stabilized by homeostatic modulation of neurotransmitter release^4^ and neurotransmitter receptors^5^. Despite considerable progress in identifying mechanisms underlying homeostatic regulation of synaptic function^4^, comparably little is known about how the molecular organization of synapses is regulated during homeostatic plasticity.

Stable neural function is especially surprising in light of a remarkable degree of ‘sub-synaptic’ molecular organization of synapses: several proteins locally enrich in ‘sub-synaptic clusters’ within presynaptic active zones (AZs)^2,6,7^, the synaptic cleft^8,9^, and the postsynaptic density (PSD)^10,11^. Moreover, sub-synaptic clusters of presynaptic proteins may align with postsynaptic clusters, including neurotransmitter receptor clusters^8,12–14^. These observations gave rise to the concept of ‘transsynaptic nanocolumns’^8,12,13^. There is some evidence that synaptic transmission predominantly occurs within transsynaptic nanocolumns^8^, suggesting that individual synapses may harbor sub-synaptic transmission channels. How transsynaptic nanocolumns are arranged within individual synapses, and how they are regulated during homeostatic plasticity remains elusive.

Several studies discovered that sub-synaptic clusters are not randomly distributed within the synaptic compartments^2,12,15^. For instance, clusters of several presynaptic proteins, including Bruchpilot (‘Brp’; CAST/ELKS)^16^, RIM-binding protein^2^, and Unc13^7^ are organized in stereotypic ring-like arrays within active zones of the *Drosophila* neuromuscular junction (NMJ). These rings are arranged in a key-lock-like fashion at specific distances from a cluster of voltagegated Ca^2+^ channels at the active zone center^2,17,18^. This stereotypic topography is thought to specify distinct functional properties of several release sites demarked by Unc13 that are driven by a common Ca^2+^-channel cluster^7^. Perturbations of this organization were shown to have profound effects on synaptic transmission and animal behavior^2,7,13,17^. Furthermore, recent data suggest the modulation of sub-synaptic active-zone organization during presynaptic homeostatic plasticity (PHP) induced by GluR impairment at this synapse^18,19^. Nevertheless, it is unclear how the specific organization of presynaptic active zones relates to postsynaptic architecture, and if homeostatic plasticity involves coordinated modulation of synaptic nano-organization across the synaptic compartments.

Compared to the specific nano-organization of presynaptic active zones, knowledge of a corresponding postsynaptic organization is scant. There is evidence for a non-homogenous and segregated distribution of AMPA and NMDA receptor clusters with regard to the PSD center of mammalian central nervous system (CNS) synapses^10,11^. Recent findings also imply a specific, ring-shaped GluR nano-organization at the *Drosophila* NMJ^20^. How postsynaptic nanoorganization relates to presynaptic architecture remains enigmatic. Moreover, it is largely unknown if postsynaptic nanoarchitecture is modulated during homeostatic plasticity.

## Results

### Transsynaptic nanocolumn rings at the *Drosophila* NMJ

Here, we explore subsynaptic molecular organization at the *Drosophila* NMJ employing dual-color stimulated emission depletion microscopy with time-gated fluorescence detection (gSTED) yielding an effective lateral resolution of < 40 nm after image deconvolution (Figure S1; see Material and Methods,^21^). To study transsynaptic organization at the nanometer scale, we imaged the presynaptic active-zone protein Brp together with postsynaptic GluRs (Figure 1A). At the *Drosophila* NMJ, Brp C-termini form sub-synaptic ring patterns at STED resolution when oriented in parallel to the imaging plane^16,22^ (Figure 1B and B’), thereby providing a proxy for synapse orientation. While confocal data did not suggest any specific GluR distribution opposite to presynaptic active zones (Figure 1A^23^), gSTED imaging revealed a distinct distribution of antibodies detecting the essential GluR subunit GluRIIC (Figure 1B and B’). In particular, we observed discrete anti-GluRIIC ‘spots’ (gray open arrowhead; Figure 1B’). A substantial fraction of GluRIIC spots appeared as ring-like patterns (white arrowheads; Figure 1B’), similar to recent observations^20^. Intriguingly, these GluRIIC rings were located in close proximity to presynaptic Brp rings (filled arrowheads; Figure 1B’). To analyze GluR distribution and its relationship to Brp, we developed an algorithm for automated ring detection (see Material and Methods and Figure S2). When detecting Brp- and GluR rings independently, the probability of detecting a GluR ring within ≤ 40 nm to a corresponding Brp-ring center was 0.84 (*n* = 89 rings; Figure S2D – E). We also noted a considerable fraction (22 ± 1%, *N* = 16 NMJs) of ring-shaped GluR fluorescence that was not apposed by presynaptic Brp fluorescence (white open arrowheads; Figure 1B’ and Figure S2E). Furthermore, some anti-GluR spots neither aligned with Brp nor formed obvious patterns (gray open arrowhead; Figure 1B’), thereby partially masking the GluR rings.

**Fig. 1.**
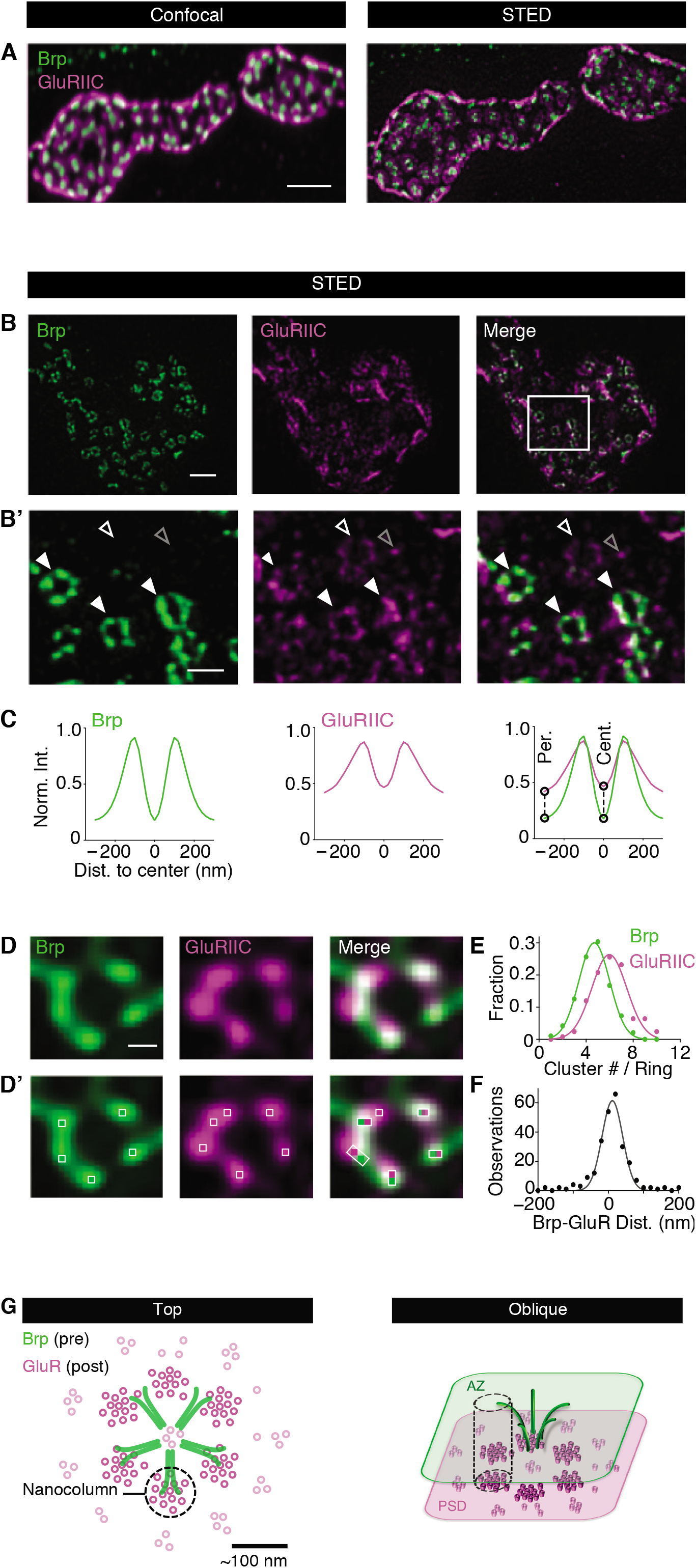
Transsynaptic nanocolumn rings at the *Drosophila* NMJ. (**A**) Maximum intensity projection of synaptic boutons of a wild-type *Drosophila* NMJ stained with anti-Brp (nc82, green); (*1*)) and anti-GluRIIC (magenta) at confocal (left) and gSTED resolution (right). (**B** and **B’**) Synaptic bouton (top) and a region within it (bottom) stained with anti-Brp (left) and anti-GluRIIC (right). Full and empty white arrowheads indicate GluR rings apposed/not apposed to presynaptic Brp rings, respectively. Empty grey arrowhead indicates ambient GluR fluorescence. (**C**) Average Brp- and GluR fluorescence intensity line profiles normalized to the respective peak obtained after Brp ring detection and subsequent analysis of the Brp- and the GluR channel (see Material and Methods). (**D and D’**) Higher magnification of Brp- (green) and GluR rings (magenta) with clusters within rings (bright pixels and white boxes, *bottom*). (**E**) Normalized histogram of local maxima number per Brp- and GluRIIC ring (“Cluster #/ Ring”) and Gaussian fit. (**F**) Histogram of nearest-neighbor distance between Brp- and GluR clusters (see Material and Methods). (**G**) Model of sub-synaptic GluR organization and Brp-GluRIIC transsynaptic alignment from top (left) and oblique (right) perspective. Scale bars: (A) 1 μm; (B) 1 μm; (B’) 200 nm; (D) 100 nm.

We next analyzed the dimensions and transsynaptic alignment of Brp- and GluR rings by anchoring line profiles in Brp-ring centers and quantifying the normalized fluorescence intensity of both channels (see Material and Methods; Figure S2D). Remarkably, there was no offset between the Brp- and GluRIIC-line profiles, indicating tight transsynaptic alignment (Figure 1C, right). Quantifying the inter-peak distance of the line profiles revealed similar ring dimensions, with slightly smaller Brp-than GluRIIC rings (Brp: 212 ± 5 nm, *n* = 703; GluRIIC: 244 ± 8 nm, *n* = 500; *P* = 0.001; Figure 1C). We also noticed significantly higher relative fluorescence intensities at the ring periphery and center in the GluRIIC-vs. the Brp-channel, as quantified by comparing relative line-profile intensities at 300 nm and 0 nm distance to the ring center (Figure 1C, right, dashed lines; both *P* < 0.001), suggesting higher background fluorescence and/or the existence of ‘ambient’ GluRs in the ring periphery and center (see below; Figure 2 and 3). These data demonstrate transsynaptic alignment between Brp- and GluR rings of similar dimensions, which are partially masked by anti-GluR fluorescence outside the rings.

**Fig. 2.**
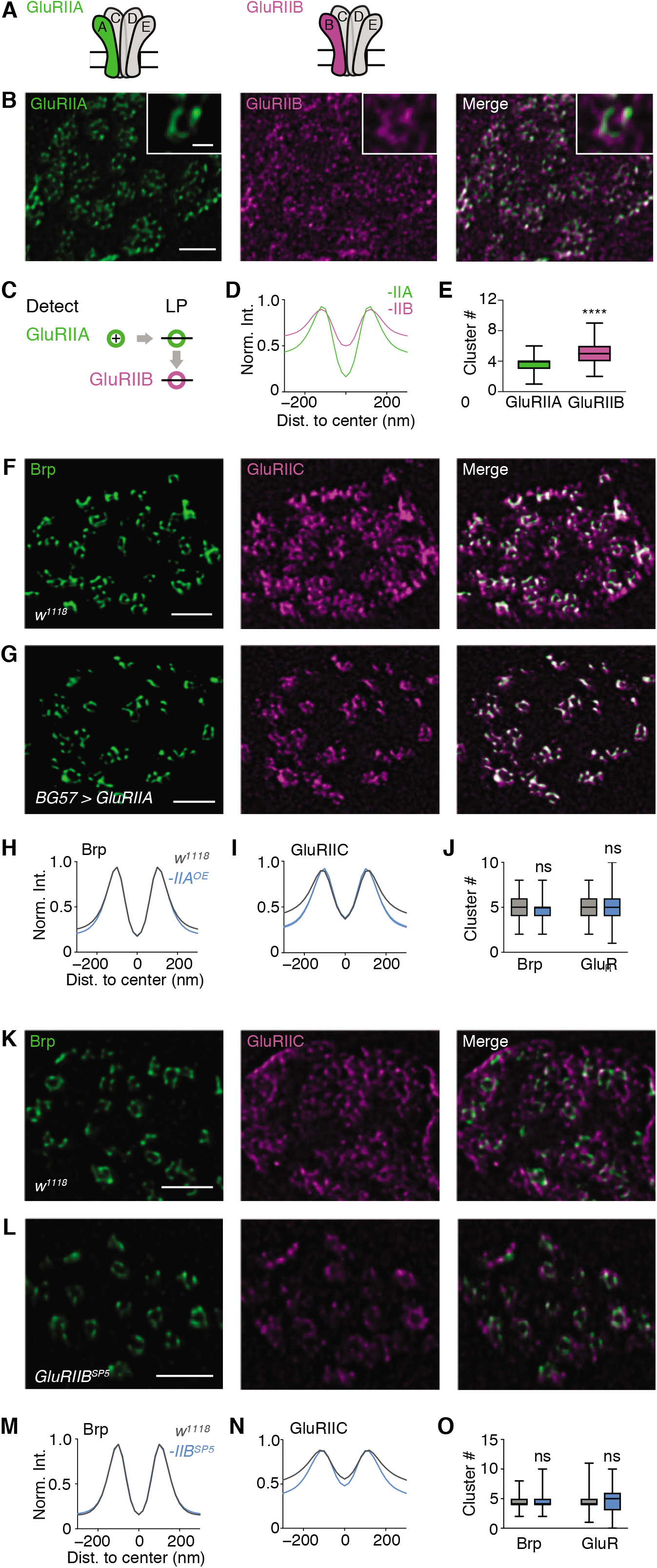
GluR-subtype specific nano-organization. (**A**) Schematic of *Drosophila* GluRIIA and GluRIIB receptor types. (**B**) Representative anti-GluRIIA (green) and GluRIIB (magenta) staining of a wildtype synaptic bouton; inset: higher magnification of a GluRIIA (left) and apposed GluRIIB (right) ring. (**C**) Schematic of GluRIIA ring detection and apposed GluRIIB rings. (**D-E**) Normalized fluorescence intensity line profiles of the GluRIIA and the GluRIIB channel (the shaded area represents s.e.m.) and average cluster number (median ±min./max.) within GluRIIA-, GluRIIB rings. GluRIIA diameter: 243 ± 7 nm, *n* = 204; GluRIIB diameter: 266 ± 7nm, *n* = 165; *P* = 0.0054. (**F-G**) Representative example boutons of *w1118* (top) and *BG57 > GluRIIA* (bottom) stained with anti-Brp (green) and anti-GluRIIC (magenta).(**H-J**) Normalized intensity line profiles and average cluster number of Brp and GluRIIC in *w^1118^* (grey) and *BG57 > GluRIIA* (blue). GluRIIC diameter: *w1118*: 248 ± 5 nm, *n* = 268; *BG57-Gal4 > UAS-GluRIIA*: 222 ± 3 nm, *n* = 257; *P* < 0.0001; Brp diameter: w^*1118*^: 222 ± 3 nm, *n* = 313; *BG57-Gal4 > UAS-GluRIIA*: 217 ± 3 nm, *n* = 282; *P* < 0.0001. (**K-L**) Representative example boutons of *w^1118^* (top) and *GluRIIB*^SP5^ (bottom) stained with anti-Brp (green) and anti-GluRIIC (magenta). (**M-O**) Normalized intensity line profiles and average cluster number of Brp and GluRIIC in w^*1118*^ (grey) and *GluRIIB*^SP5^ (blue). GluRIIC diameter: *w^1118^*: 265± 5 nm, *n =* 274; *GluRIIBsp5*: 239 ± 4 nm, *n* = 442; *P* < 0.0001; diameter Brp: w^*1118*^: 214 ± 1.9 nm, *n* = 374; *GluRIIB^sp5^:* 211 ± 2 nm, *n* = 573; *P* = 0.26. Scale bars: (B, top) 200 nm;(B, bottom) 1 μm; (F) 1 μm; (G) 1 μm; (K) 1 μm; (L) 1 μm.

Within Brp- and GluR rings, fluorescence intensity was heterogeneously distributed (Figure 1D). Earlier work revealed ~5-6 local fluorescence intensity maxima per Brp ring at STED resolution, implying that Brp rings are composed of ~5-6 distinct Brp ‘clusters’^24^ (Figure 1D and D’). To analyze fluorescence within the rings, we developed an algorithm for local fluorescence intensity maximum detection (see Material and Methods; Figure 1D’ and S3). This analysis gave an average number of 4.7 ± 0.1 local maxima per Brp ring (*n* = 191; Figure 1D’, white squares; Figure 1E), and 6.0 ± 0.1 local maxima per GluRIIC ring (*n* = 125; Figure 1D’ and 1E). Previous direct stochastic optical reconstruction microscopy (dSTORM) data suggest that the local maxima within the Brp ring correspond to Brp clusters/filament bundles, each consisting of ~30 Brp molecules^25^. Correspondingly, the local maxima in the GluR channel likely reflect GluR clusters, similar to findings at mammalian synapses^10,11,26^. These observations are consistent with the idea that Brp- and GluR rings are composed of ~5 and ~6 clusters, respectively.

We next explored the relationship between Brp- and GluR fluorescence intensity maxima, henceforth called ‘clusters’, within the rings by quantifying their lateral nearest-neighbor distance. This analysis gave an average lateral nearest-neighbor distance of 14 ± 5 nm (*n* = 256) between Brp- and GluRIIC clusters (Figure 1F), below our lateral resolution (Figure S1). Line-profile analysis along the Brp ring circumference revealed that 77 ± 3.4 % of the Brp clusters tightly align with GluRIIC clusters (n=30; see also Figure S3C), suggesting that most Brp clusters are apposed by a GluR cluster. In addition, we observed unaligned GluRIIC clusters within the rings (Figure 1D and S3C). Taken together, our experiments uncovered transsynaptically aligned Brp-GluR rings composed of ~4-5 tightly aligned Brp- and GluR clusters, as well as ~1-2 unaligned GluR clusters. Previous studies termed transsynaptically-aligned synaptic protein clusters ‘transsynaptic nanocolumns’^8,13^. Based on this terminology, our data support a model of transsynaptic nanocolumn rings (Figure 1G). Previous work demonstrated that a Ca^2+^-channel cluster localizes to the Brp ring center^7,24^. Thus, the stereotypic ring topography likely reflects the organization of transsynaptic nanocolumns around a central Ca^2+^-channel cluster, which triggers release in response to synaptic stimulation (see Discussion).

### GluR subunit manipulations unmask transsynaptic nanocolumn rings

Transsynaptic Brp-GluR nanocolumn rings are partially masked by anti-GluR fluorescence outside the rings (Figure 1B’, open grey arrowhead). We next aimed at unmasking transsynaptic nanocolumn rings and investigating GluR nano-organization. *Drosophila* GluRs are heterotetramers composed of three essential subunits (GluRIIC, GluRIID and GluRIIE) and either a GluRIIA- or a GluRIIB subunit that determine receptor desensitization^27,28^ (Figure 2A). We therefore hypothesized that sub-synaptic GluR organization may be GluR-subtype specific and investigated GluRIIA and GluRIIB distribution with STED microscopy. Analysis of anti-GluRIIA and anti-GluRIIB co-stainings revealed ringlike arrays of GluRIIA- and GluRIIB fluorescence of similar dimensions (Figure 2B – D), suggesting that receptors within the rings incorporate both, the GluRIIA and the GluRIIB subunit. GluRIIA-centered line profile analysis showed significantly higher relative GluRIIB fluorescence intensities towards the ring periphery and in the ring center compared to GluRIIA (GluRIIA: *n* = 204; GluRIIB: *n* = 165; *P* < 0.0001; Figure 2D), either indicating that ambient GluRs outside the rings predominately contain GluRIIB, or anti-GluRIIB background fluorescence (see below). Within the rings, we detected ~4 or ~5 clusters in the GluRIIA- and the GluRIIB channel, respectively (*P* < 0.0001; Figure 2E). As individual rings contain on average ~5 Brp clusters and ~6 GluRIIC clusters (Figure 1E), this implies that most transsynaptic nanocolumns likely harbor both, GluRIIA- and GluRIIB-containing receptors.

While anti-GluRIIA fluorescence was largely confined to the rings, anti-GluRIIB fluorescence was found inside and outside the rings (Figure 2B, D). To test whether the differential anti-GluRIIA and anti-GluRIIB fluorescence distribution is due to a GluR subtype-specific nano-organization, we assayed GluR organization after genetic manipulation of these two subunits. GluRIIA-containing GluRs primarily localize to rings (Figure 2B and D). We thus hypothesized that GluR rings are more distinct upon GluRIIA overexpression. GluRIIC fluorescence indeed appeared more distinct and ring-like after postsynaptic *GluRIIA* overexpression (*BG57-Gal4 > UAS-GluRIIA*) compared to wild-type controls (Figure 2F and G, *middle*). Line profile analysis of aligned Brp-GluRIIC rings revealed significantly lower relative GluRIIC fluorescence intensity at the ring periphery, but not in the ring center, upon GluRIIA overexpression (*w^1118^*: *n* = 268; *BG57-Gal4 > UAS-GluRIIA: n* = 257; periphery: *P* < 0.0001; center: *P* > 0.99; Figure 2I), implying more distinct GluR rings due to reduced GluR density in the ring periphery.

*GluRIIA* overexpression also resulted in a small, but significant reduction of relative Brp-fluorescence intensity at the ring periphery (w^*1118*^: *n* = 313; *BG57-Gal4 > UAS-GluRIIA: n* = 282; *P* < 0.0001; Figure 2H), indicating a transsynaptic change. Intriguingly, while GluRIIC rings were slightly broader than Brp rings in wild type (Figure 1C, Figure S4), there was no offset between the peaks of GluRIIC- and Brp line profiles after GluRIIA overexpression, suggesting tight transsynaptic alignment between the rings (Figure S4A and B). We did not observe significant changes in cluster number within Brp- and GluRIIC rings between the two genotypes (Brp: *P* = 0.26; GluRIIC: *P* = 0.77; Figure 2J), implying that GluRIIA overexpression does not affect transsynaptic nanocolumn number. Thus, GluRIIA predominately localizes to transsynaptic nanocolumn rings and GluRIIA overexpression unmasks tightly aligned transsynaptic nanocolumn rings.

Anti-GluRIIB fluorescence outside the rings indicates that ambient receptors may preferentially incorporate the GluRIIB subunit (Figure 2B and D). Loss of the GluRIIB subunit is therefore expected to decrease anti-GluRIIC levels outside the rings, and to result in more distinct GluR rings. We therefore used CRISPR/Cas9 targeted mutagenesis to generate null mutations in the *GluRIIB* subunit (Materials and Methods). *GluRIIB^SP5^* mutants displayed distinct GluRIIC rings (Figure 2K and L), and relative GluRIIC-line profile intensity was significantly dimmer towards the ring periphery and in the ring center in *GluRIIB^SP5^* mutants compared to wild type (*w^1118^*: *n* = 274; *GluRIIB^SP5^*: *n* = 442; periphery: *P* < 0.0001; center: *P* = 0.001; Figure 2N). Brp line profiles (Figure 2M), as well as Brp-and GluRIIC cluster number were similar between *GluRIIB^SP5^* and wild type (line profile: periphery: *P* = 0.84; center: *P* > 0.99; cluster: *w^1118^*: *n* = 374; *GluRIIB^SP5^*: *n* = 573; Brp: *P* = 0.91; GluRIIC: *P* = 0.99; Figure 2M-O), indicating that loss of the GluRIIB-subunit does not affect Brp dimensions and Brp-/GluRIIC nanocolumn number. We conclude that GluRIIB-containing receptors localize within and outside of transsynaptic nanocolumn rings. Moreover, *GluRIIA* overexpression and *GluRIIB* loss unmask transsynaptic nanocolumn rings, and support a model of GluR subtype-specific nano-organization.

Previous work revealed no defects in spontaneous or AP-evoked synaptic transmission upon *GluRIIA* overexpression^28^ or after *GluRIIB* loss^23^. Moreover, we show that mEPSP amplitudes are even increased in *GluRIIB^SP5^* mutants compared to controls (Figure S4C). Thus, two genetic manipulations that mainly decrease ambient receptors, but not receptors inside the nanocolumn ring, do not induce a corresponding decrease in synaptic transmission, indicating that synaptic transmission is largely confined to transsynaptic nanocolumn rings (see Discussion).

### Transsynaptically-aligned Neto-β rings stabilize GluRs

We next sought to provide independent evidence for transsynaptically-aligned rings by analyzing the sub-synaptic distribution of Neto, an auxiliary GluR subunit previously suggested to play a role in GluR clustering^29,30^ (Figure 3A). There are two Neto isoforms with different expression patterns at the *Drosophila* NMJ: while Neto-α is expressed both, pre- and postsynaptically^31^, Neto-β is the major postsynaptic isoform at the *Drosophila* NMJ^30^. Anti Neto-β arranged in ring-like arrays in close proximity to presynaptic Brp (Figure 3B and B’). We also observed a significant fraction of anti-Neto-β that was not apposed by Brp, similar to GluRIIC (Figure 1B and B’). Line-profile analysis revealed an overlap between Brp- and Neto-β-line profile peaks (Figure 3D), demonstrating similar dimensions and transsynaptic alignment. Similar to GluRIIC, Neto-β fluorescence intensity was higher at the ring periphery and center compared to Brp (Neto-β: *n* = 149; Brp: *n* = 274; periphery: *P* < 0.0001; center: *P* < 0.0001; Figure 3D), suggesting that Neto-β not only localizes opposite to Brp C-termini. Neto-β line profiles also aligned with anti-GluRIIA line profiles (Figure S4A-B). The close relationship between Neto-β and GluRIIA rings is in line with genetic data suggesting that Neto-β predominantly stabilizes GluRIIA-containing receptors^30^. However, the Neto-β fluorescence outside the rings (Figure 3B’ and D) indicates that this auxiliary GluR subunit also interacts with other proteins, or the existence of a Neto-β reserve pool. Within transsynaptically-aligned Neto-β rings, we detected 3.3 ± 1.5 clusters (*n* = 397; Figure 3E), significantly fewer than for Brp (~5; Figure 1E; *P* < 0.0001) or GluRIIC (~6; GluRIIA vs. Neto-β cluster number: Figure S5C; *P* < 0.0001), suggesting that Neto-β is unlikely part of every nanocolumn. We conclude that Neto-β forms rings that are transsynaptically aligned with Brp, and which are composed of ~3 clusters.

**Fig. 3.**
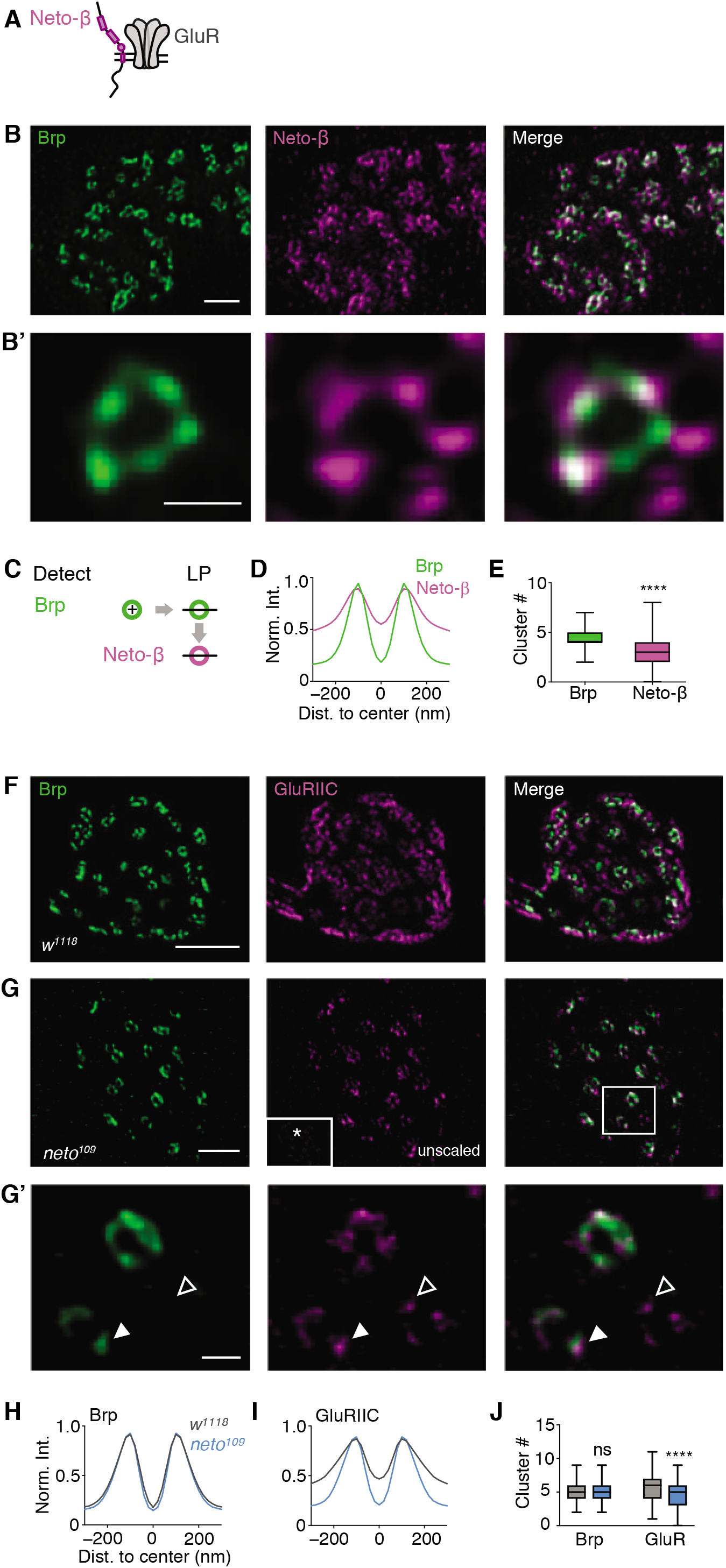
Transsynaptically-aligned Neto-β rings stabilize GluRs. (**A**) Schematic of Neto-β and GluRs at the *Drosophila* NMJ. (**B**) Representative anti-Brp (green) and anti-Neto-β staining of a wild-type synaptic bouton. (**B’**) Higher magnification of a Brp (left) and corresponding Neto-β (right) ring. (**C**) Schematic of Brp ring detection and apposed Neto-β ring. (**D**) Normalized fluorescence intensity line profiles of the Brp and the Neto-β channel. (**E)** Average cluster number (median±min./max.) within Brp- and Neto-βrings. Brp diameter: 209.2 ± 2nm, *n* = 397; Neto-β diameter: 241 ± 4 nm, *n* = 292; *P* < 0.0001. (**F-G’**) Representative example boutons of *w^1118^* (top) and *neto^109^* (bottom) stained with anti-Brp (green) and anti-GluRIIC (magenta). (**G**, middle) *neto^109^* GluRIIC intensity scaled to *w^1118^* is indicated in the white box (*).(**G’**) Higher magnification of *neto^109^* Brp- (left) and apposed GluRIIC (right) rings. The filled and open arrowhead highlight transsynaptically aligned and unaligned GluR clusters, respectively.(**H-J**) Normalized intensity line profiles and average cluster number of Brp and GluRIIC in *w^1118^* (grey) and *neto*^109^ (blue). GluRIIC diameter: *w^1118^*: 245 ± 9 nm, *n* = 103; *neto*^109^: 217 ± 4 nm, *n* = 143; *P* = 0.02; Brp diameter: *w^1118^*: 216± 4nm, *n* = 133; *neto*^109^: 217 ± 3 nm, *n* = 149; *P* = 0.77. Scale bars: (B) 1 μm; (B’) 200 nm; (E) 1 μm; (F) 1 μm; (F’) 200 nm.

Neto is thought to be required for GluR clustering at the *Drosophila* NMJ^29^. We next assessed how loss of *neto* affects sub-synaptic organization. GluRIIC fluorescence intensity was strongly decreased in hypomorphic *neto^109^* mutants, in which Neto-α and Neto-β levels are strongly reduced^29^ at confocal resolution (Figure S5D-E), implying GluR loss, in line with previous work^29^. At gSTED resolution, GluRIIC rings appeared dimmer, and GluRIIC intensity outside the rings was markedly reduced (Figure 3G, middle, inset), suggesting GluR loss inside and outside the rings. As evident from the bouton with scaled GluRIIC fluorescence intensity shown in Figure 3G (middle), the remaining GluRs formed very distinct rings that transsynaptically aligned with Brp rings (Figure 3G - G’). We observed a significant increase in the fraction of ‘ring-like GluRIIC line profiles’ in *neto^109^* mutants (Figure S5F), and a significant decrease in normalized fluorescence intensity in the ring periphery and center in *neto^109^* mutants (*w^1118^*: *n* = 103; *neto^109^*: *n* = 143; periphery and center: *P* < 0.0001; Figure 3I), indicating that GluR levels outside the ring were decreased more strongly than inside the ring in *neto^109^* mutants. Together with the decrease in ambient GluR levels upon *GluRIIA* overexpression or in *GluRIIB* mutants (Figure 2), these data again imply that GluRs reside at the ring periphery and center of wild-type synapses (see also Figure S3). We also noted a slight, but significant decrease in GluRIIC cluster number within the rings of *neto^109^* mutants (*w^1118^*: *n* = 133; *neto^109^*: *n* = 149; *P* < 0.0001; Figure 3J). The decreased GluR levels in *neto^109^* mutants also led to more distinct GluR clusters within the rings (Figure 3G’, middle, arrowheads), likely caused by a lower GluR abundance per cluster. While we observed some GluR clusters within the rings that were not apposed by Brp fluorescence (Figure 3G’, middle, open arrowhead), most GluR clusters localized in close proximity to Brp (Figure 3G’, middle, filled arrowhead), indicating a tight transsynaptic alignment, thereby further supporting a model of transsynaptic nanocolumn rings^8,12^. We did not observe apparent changes in Brp intensity (periphery: *P* = 0.196; center: *P* = 0.22), or Brp-cluster number between the two genotypes (*w^1118^*: *n* = 133; *neto^109^*: *n* = 149; *P* ≥ 0.99; Figure 3H-J). Taken together, the investigation of the auxiliary GluR subunit Neto provides independent evidence for postsynaptic ring patterns that are aligned across the synaptic cleft and revealed that Neto stabilizes GluRs inside and outside the nanocolumn rings. Furthermore, reduced Neto levels resulted in a remarkable degree of alignment between Brp- and GluRIIC clusters within the rings, consistent with a model of transsynaptic nanocolumns.

### Rapid homeostatic modulation of transsynaptic nanocolumn rings

Having established the presence of transsynaptic nanocolumn rings, we next asked if they are modulated during synaptic plasticity. At the *Drosophila* NMJ, pharmacological GluR impairment induces a homeostatic increase in neurotransmitter release that precisely compensates for the perturbation within minutes after receptor impairment^4,32–34^. There is evidence that Brp content per AZ and Brp cluster number is increased within ten minutes during this form of homeostatic plasticity, commonly referred to as ‘presynaptic homeostatic plasticity’ (PHP)^19^. Based on the transsynaptic alignment between Brp- and GluR rings, we hypothesized a modulation of transsynaptic Brp-GluR nanocolumns during PHP. First, we employed confocal microscopy to probe relative changes in Brp- and GluR fluorescence intensity during homeostatic plasticity (Figure 4A-E). Application of the GluR antagonist philanthotoxin-433 (PhTX) for 5’ decreased mean Brp fluorescence intensity compared to saline-treated controls (HL3-saline: *n* = 756; PhTX: *n* = 940; - 8.0%; *P* < 0.0001; Figure 4A, B, C and E), implying a slight decrease in Brp levels. The decrease in Brp intensity was also accompanied by a significant decrease in Brp area (Figure S6A), suggesting Brp redistribution after 5’ PhTX incubation. By contrast, we noted a prominent increase in GluRIIC-fluorescence intensity after PhTX treatment for 5’ (HL3-saline: *n* = 777; PhTX: *n* = 1066; +23.0%; *P* < 0.0001; Figure 4A, B, D and E), suggesting increased GluR abundance. Furthermore, GluR area significantly decreased after 5’ PhTX application (Figure S6A), implying that the increase in GluR intensity may be due to GluR redistribution (Figure S6A). After 15’ of PhTX treatment, there was a significant increase in both, Brp-fluorescence intensity (HL3: *n* = 1126; PhTX: *n* = 844; +25.6%; *P* < 0.0001; Figure 4E), and GluRIIC-fluorescence intensity compared to controls (HL3: *n* = 1280; PhTX: *n* = 724; +31.8%; < 0.0001; Figure 4E). Similar results were obtained after PhTX incubation for 30’ (Brp: HL3: *n* = 1910; PhTX: *n* = 1736; +5%; *P* < 0.0001; GluRIIC: HL3: *n* = 1386; PhTX: *n* = 1333; +16%; *P* < 0.0001; Figure 4E). These data provide evidence that pharmacological GluR perturbation induces a rapid and sequential increase in synaptic GluRIIC and Brp abundance on the minute time scale, with GluRIIC modulation preceding Brp modulation at confocal resolution.

**Fig. 4.**
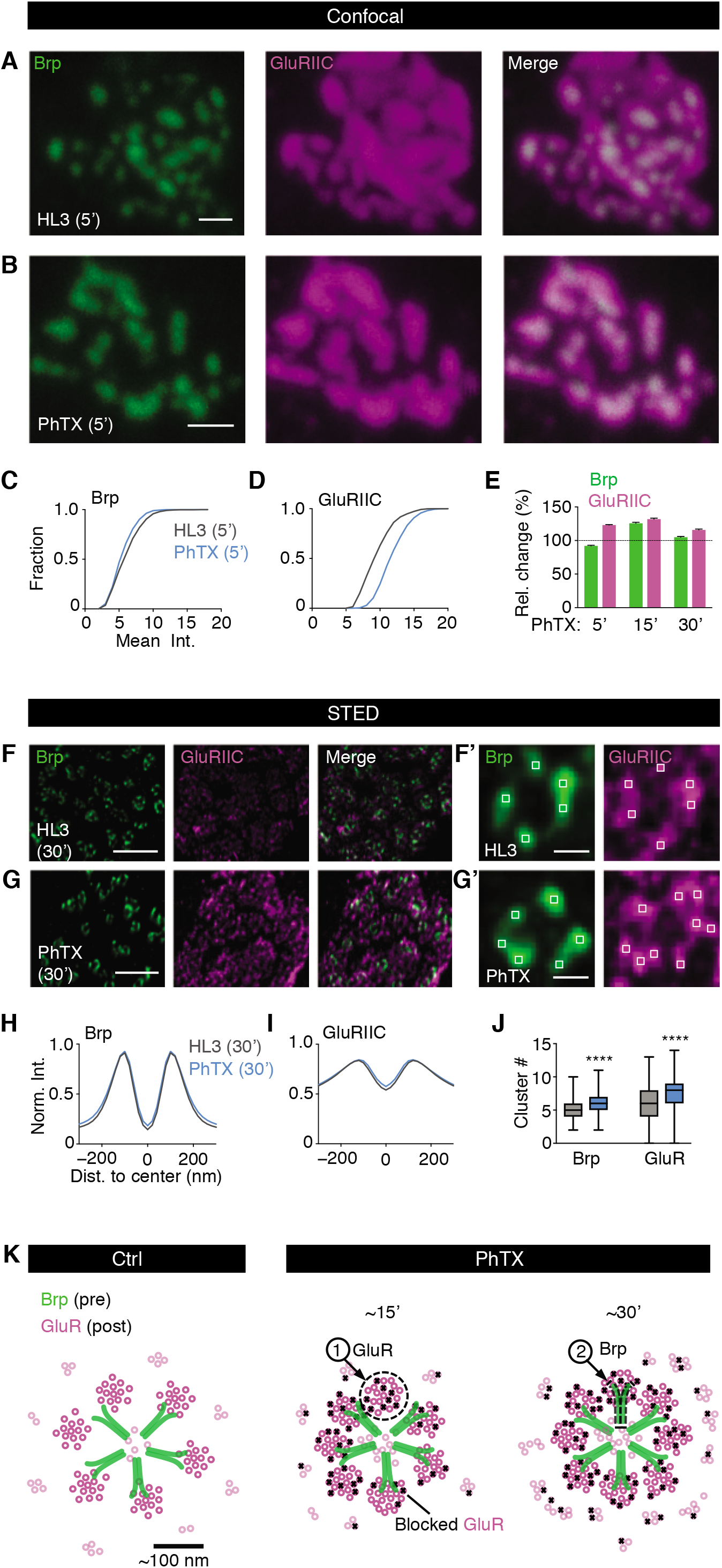
Rapid and sequential modulation of transsynaptic nanocolumn rings during homeostatic plasticity. (**A, B**) Representative boutons stained with anti-Brp and anti-GluRIIC upon HL3/ saline or PhTX treatment (5’) in *w^1118^* at confocal resolution. (**C, D**) Corresponding cumulative frequency histogram of mean intensity changes of the Brp (left) and the GluRIIC (right) channel upon HL3/saline (grey) or PhTX (blue) (5’). (**E**) Relative change of the Brp and GluRIIC mean intensity upon PhTX incubation for 5’, 15’ and 30’. (**F, G**) Representative boutons stained with anti-Brp and anti-GluRIIC upon HL3 (top) or PhTX treatment (30’) in *w^1118^* at gSTED resolution. (**F’, G’**) Brp and GluRIIC ring examples with cluster number/ring indicated by white boxes.(**H, I**) Corresponding normalized intensity line profiles of the Brp and GluRIIC channel upon HL3 (grey) or PhTX (blue) treatment (both 30’). (**J**) Average cluster number within Brp (green) and GluRIIC (magenta) rings upon HL3 or PhTX treatment. GluRIIC diameter: HL3: 296 ± 5 nm, n = 500; PhTX: 281 ± 5 nm, n = 487; *P* = 0.04; Brp diameter: HL3: 217 ± 2, n = 703; PhTX: 215 ± 2, n = 705; *P* = 0.57. (**K**) Model showing changes in sub-synaptic organization of transsynaptically aligned Brp and GluR rings upon PhTX-induced GluR perturbation. Note that the model does not reflect changes seen at confocal resolution (Fig. S6). Scale bars: (A) 1 μm; (B): 1 μm; (F) 1 μm; (F’) 100 nm; (G) 1 μm; (G’) 100 nm, (K) 100 nm.

We next used STED imaging to investigate potential changes in transsynaptic architecture during homeostatic plasticity (Figure 4F-J). While we did not detect significant changes in the organization of Brp- and GluR rings after PhTX application for 5’ (Figure S6C), there was a significant increase in GluR cluster number (HL3: *n* = 500; PhTX: *n*=487; *P* = 0.008), but no significant change in Brp-cluster number (*P =* 0.80) upon PhTX incubation for 15’ (Figure S6C). After PhTX exposure for 30’, both GluRIIC- and Brp cluster number significantly increased (HL3: *n* = 703; PhTX: *n* = 705; both *P* < 0.0001; Figure 4F’-G’, J and Figure S6C), without major changes in GluRIIC- and Brp-ring diameter (GluRIIC: HL3: *n* = 500; PhTX: *n* = 487; *P* = 0.04; Brp: HL3: *n* = 703; PhTX: *n* = 705; *P* = 0.57; Figure 4H-I). These observations suggest transsynaptically-coordinated modulation of synaptic nano-architecture during PHP. Consistent with our confocal data, the increase in GluR-cluster number precedes the increase in Brp-cluster number, implying sequential modulation of GluR- and Brp-cluster number during PHP. The delay between the changes at confocal and STED resolution may indicate that changes in GluR- and Brp-distribution and/or levels precede the increase in nanocolumn number (see Figure S6). Together, our data suggest rapid, sequential modulation of transsynaptic Brp-GluR nanocolumn rings during homeostatic plasticity (Figure 4K).

### Homeostatic modulation of transsynaptic nano-oganization and presynaptic release requires *neto*

Based on our findings that (*i*) Neto-β aligns with GluRIIA and Brp, (*ii*) Neto stabilizes GluRs inside and outside the rings, and that (*iii*) both, GluRs and Brp undergo rapid changes during homeostatic plasticity, we hypothesized that Neto-β is modulated during homeostatic plasticity. Application of PhTX to wild-type synapses for 15’ resulted in a significant increase in the mean Neto-β fluorescence intensity at confocal resolution compared to saline-treated controls (HL3: *n* = 3278; PhTX: *n* = 772; *P* < 0.0001; Figure 5A-C). Additionally, we detected a significant increase in Neto-β cluster number per ring upon PhTX treatment for 15’ using STED imaging (HL3: *n* = 268; PhTX: *n*= 237; *P* < 0.0001; Figure 5D), suggesting the modulation of nano-cluster abundance of the auxiliary GluR subunit Neto-β during homeostatic plasticity. These data provide independent evidence for the modulation of postsynaptic nano-organization during homeostatic plasticity.

**Fig. 5.**
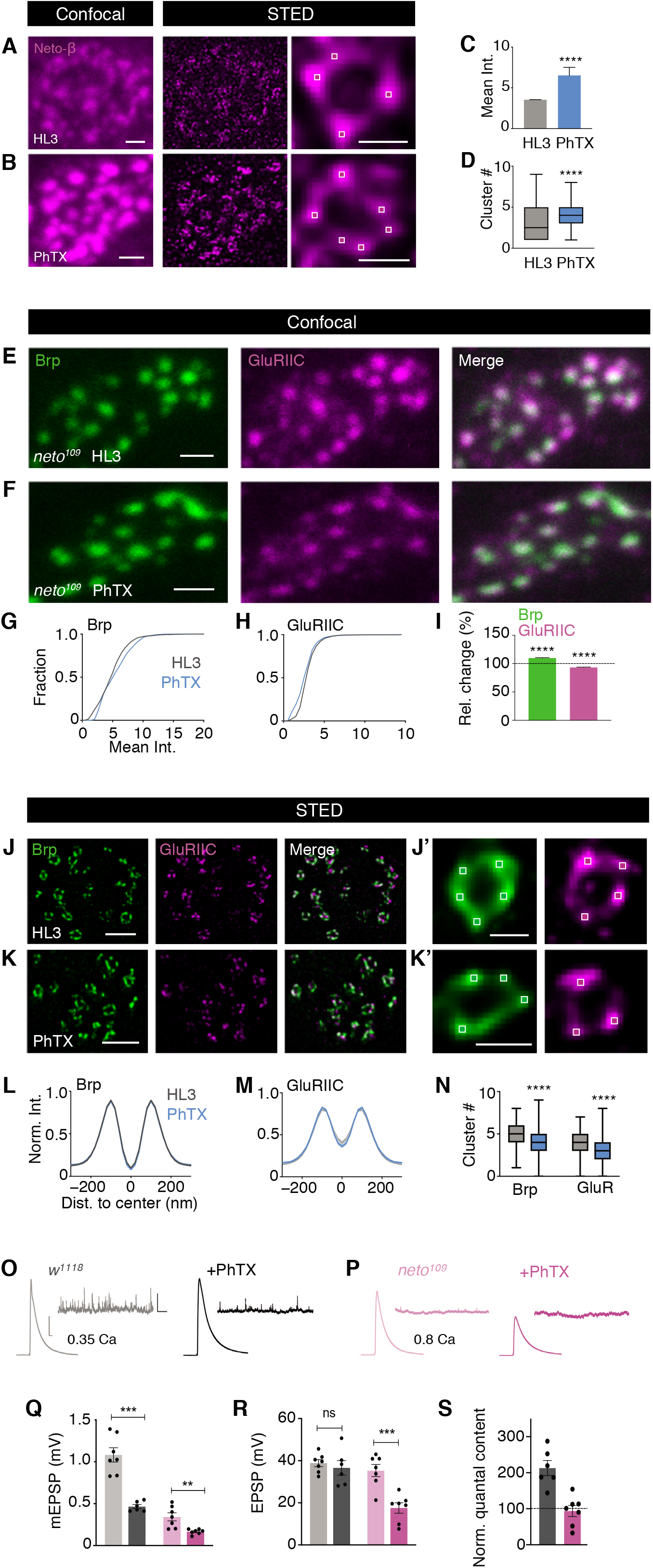
Rapid homeostatic increase in GluR abundance and transsynaptic nanocolumn number requires neto. (**A-B**) Representative boutons stained with anti-Neto-β upon HL3/saline or PhTX treatment (15’) in *w^1118^* at confocal (left) and STED (middle) resolution and higher magnification of Neto-β ring (left). (**C-D**) Corresponding mean intensity and cluster number changes of Neto-β upon HL3/saline (grey) or PhTX (blue) (15’). (**E, F**) Representative *neto^109^* boutons stained with anti-Brp and anti-GluRIIC upon HL3 or PhTX treatment (15’) at confocal resolution. (**G-I**) Corresponding cumulative frequency histogram of mean intensity changes and relative changes of the Brp (left) and the GluRIIC (right) channel upon HL3/ saline (grey) or PhTX (blue) in *neto^109^*. (**J-K**) Representative boutons stained with anti-Brp and anti-GluRIIC upon HL3 or PhTX treatment (15’) in *neto^109^* at gSTED resolution. (**J’-K’**) Brp and GluRIIC ring example incubated with HL3 or PhTX with average cluster number/ring indicated in white boxes. (**L-N**) Corresponding normalized intensity line profiles of the Brp and GluRIIC channel upon HL3 (grey) or PhTX (blue) treatment (15’) and average cluster number within Brp (green) and GluRIIC (magenta) rings upon HL3 or PhTX treatment in *neto^109^*. GluRIIC diameter: HL3: 296 ± 5 nm, n = 500; PhTX: 281 ± 5 nm, n = 487; *P* = 0.04; Brp diameter: HL3: 217 ± 2, n = 703; PhTX: 215 ±2, n = 705; *P* = 0.57. (**O**) ** Example mEPSP and EPSP traces in w1118 upon HL3 (grey) or PhTX (black) treatment. (**P**) Example mEPSP and EPSP traces in *neto^109^* upon HL3 (light pink) or PhTX (dark pink) treatment. (**Q-R**) Average mEPSP and EPSP in *w^1118^* and *neto^109^* upon HL3 or PhTX treatment. (**S**) Average normalized quantal content (QC = EPSP/mEPSP) in *w^1118^* and *neto^109^* upon PhTX treatment. Scale bars: (A, left) 1 μm; (A, right) 200 nm; (B, left): 1 μm; (B, right) 200 nm; (E) 1 μm; (F) 1 μm; (J) 500 nm; (J’) 200 nm; (K) 500 nm; (K’) 200 nm; (O) EPSP: horizontal scale: 10 ms; vertical scale: 10 mV; mEPSP: horizontal scale: 2000 ms; vertical scale: 2 mV.

Given the changes in Neto-β nanoorganization during PHP, we investigated if *neto* is required for the modulation of transsynaptic nano-organization during homeostatic plasticity. PhTX treatment for 15’, a manipulation that robustly increases GluRIIC abundance in wild type (Figure 4A-E), did not increase, but rather slightly decreased mean GluRIIC fluorescence intensity in *neto^109^* mutants at confocal resolution (HL3: *n* = 2363; PhTX: *n* = 2497; *P* < 0.0001; Figure 5E-I). Brp-fluorescence intensity was significantly increased by ~10% after PhTX application at *neto^109^* mutant synapses (HL3: *n* = 2162; PhTX: *n* = 1775; *P* < 0.0001; Figure 5E-I), but significantly less pronounced than in wild type (~23%; *P* < 0.0001; Figure 4E; see below). STED imaging revealed a slight decrease in GluRIIC- and Brp-cluster number per ring upon PhTX treatment (15’) in *neto^109^* mutants (Brp: HL3: *n* = 287; PhTX: *n* = 253; GluR: HL3: *n* = 274; PhTX: *n* = 247; both: *P* < 0.0001; Figure 5J’, K’ and N), without significant changes in GluRIIC- and Brp-ring dimensions (HL3: *n* = 332; PhTX: *n* = 295; both: *P* < 0.0001; Figure 5L-M). Hence, the rapid homeostatic increase in GluR abundance, GluR-cluster number, and transsynaptic nanocolumn number requires wild-type Neto levels.

Based on the observation that the modulation of transsynaptic nanoarchitecture during PHP is impaired in *neto^109^* mutants, we next asked if homeostatic modulation of synaptic function is affected in this genetic background. While Neto-α has been implicated in PHP presynaptically^31^, it is unknown if PHP is impaired in *neto^109^* mutants with reduced Neto-α and Neto-β levels^29,31^. PhTX application for 15’ to wildtype NMJs led to a ~50% decrease in mEPSP amplitude compared to untreated controls (HL3) (HL3: *N* = 7; PhTX: *N* = 6; *P* = 0.002; Figure 5O-Q), indicating GluR impairment. PhTX treatment did not change EPSP amplitude in wild type (HL3: *N* = 7; PhTX: *N* = 6; *P* = 0.575; Figure 5O and R), translating into a significant increase in quantal content (= EPSP/mEPSP; *N* = 6; *P* = 0.003; Figure 5S) compared to controls, suggesting increased presynaptic release, consistent with PHP^32^. *neto^109^* mutants exhibited a pronounced decrease in mEPSP amplitude in the absence of receptor perturbation (Figure 5P and Q), suggesting impaired synaptic transmission, in line with earlier work^29^. In contrast to wild type, PhTX incubation led to a similar decrease in mEPSP and EPSP amplitude in *neto^109^* mutants (*N* = 7; mEPSP: *P* = 0.009; EPSP: *P* = 0.0006; Figure 5P - R; note that recordings were conducted at elevated extracellular Ca^2+^ concentration to compensate for the defect in baseline synaptic transmission), resulting in no change in quantal content (*N* = 7; *P* = 0.69; Figure 5S). These data demonstrate impaired PHP induced by pharmacological GluR inhibition in *neto^109^* mutants. As *neto^109^* mutants also display a defect in homeostatic modulation of transsynaptic nanoarchitecture (Figure 5J - N), these data demonstrate that wild-type Neto levels are required for homeostatic control of synaptic nano-architecture and function.

## Discussion

In this study, we identified a stereotypic arrangement of transsynaptically-aligned molecular nanocolumns that is regulated in a modular and sequential fashion during homeostatic plasticity at the *Drosophila* NMJ. Moreover, we revealed a GluR subtype-specific nano-organization and discovered that the auxiliary GluR subunit Neto is essential for rapid homeostatic modulation of transsynaptic nanocolumn number and neurotransmitter release.

Previous work demonstrated that a cluster of voltage-gated Ca^2+^ channels localizes to the Brp-ring center at the *Drosophila* NMJ. Furthermore, Unc13, a molecule suggested as a molecular correlate of presynaptic release sites^6,7^, forms ring-like arrays in close proximity to Brp’s C-termini^6^. In light of these previous findings, our results are consistent with a model in which Ca^2+^ influx at the Brp-/AZ center induces neurotransmitter release in the nanocolumn rings. Given that the neurotransmitter content released by a single synaptic vesicle does not activate all GluRs of a given postsynaptic density at the *Drosophila* NMJ^35^, and that *Drosophila* GluRs have a low glutamate affinity^28^, neurotransmitter release likely mainly activates GluRs that are aligned to presynaptic release sites. The ring geometry may thus represent the organization of transsynaptic nanocolumns around a shared Ca^2+^-channel cluster. There is some evidence that synaptic transmission predominantly occurs within transsynaptic nanocolumns^8^. Hence, the transsynaptic nanocolumn rings discovered here may reflect sub-synaptic transmission modules that are activated by a common Ca^2+^-channel cluster. Future work is needed to assess how many GluRs are activated by the release from one synaptic vesicle.

GluR subunit composition and GluR location with regard to release sites are important factors determining synaptic efficacy^36^. At the *Drosophila* NMJ, the ratio of slowly and rapidly desensitizing GluRIIA- and GluRIIB-containing receptors is a key regulator of quantal size^28^. We revealed that transsynaptic nanocolumns harbor a mix of GluRIIA- and GluRIIB-containing receptors, and that ambient receptors mainly incorporate the GluRIIB subunit. Based on the increased mEPSP amplitude seen in *GluRIIB^SP5^* mutants, our data suggest that GluRIIB-containing receptors surrounding the nanocolumns have the potential to negatively regulate synaptic transmission by replacing GluRIIA-containing receptors within the nanocolumns.

A variety of auxiliary subunits controls GluR assembly, trafficking, and function^37^. The auxiliary GluR subunit Neto has been implicated in GluR clustering at the *Drosophila* NMJ^29^. We uncovered modular ring arrays of Neto-β that transsynaptically align with Brp, suggesting that this auxiliary GluR subunit is a postsynaptic element of transsynaptic nanocolumn rings. In contrast to *neto^109^* mutants - in which both, Neto-α and Neto-β levels are reduced^31^ - loss of *neto-α* does not decrease GluR levels or mEPSP amplitude^31^, suggesting that this Neto isoform does either not stabilize GluRs at the *Drosophila* NMJ, or a compensation by Neto-β. While reduced levels of ambient receptors do not impair synaptic transmission in case of *GluRIIA* overexpression^28^ or in *GluRIIB^SP5^* mutants (Figure S4C), the decreased GluR abundance within the rings of *neto^109^* mutants correlates with a decrease in spontaneous and AP-evoked synaptic transmission (Figure 5Q - R) 29, implying that synaptic transmission predominantly occurs within the rings.

GluR impairment at the *Drosophila* NMJ induces a homeostatic increase in release^32,33^, and there is evidence for the modulation of presynaptic nano-architecture during this form of homeostatic plasticity^19^. A recent study reported increased GluR levels upon sustained pharmacological GluR inhibition for several days^20^. We here demonstrate GluR modulation within minutes after pharmacological GluR impairment that precedes the modulation of Brp. Moreover, GluR- and Brp-fluorescence intensity changes seen with confocal microscopy preceded the increase in GluR- and Brp-cluster number at STED resolution. These data imply that postsynaptic modulation precedes presynaptic regulation, and implies that GluR- and Brp levels and/or distribution are modulated prior to the formation of new transsynaptic nanocolumns. The temporal sequence of GluR- and Brp regulation may also explain the existence of GluR clusters within the ring that are not apposed by Brp. Together, these findings support a model of coordinated, transsynaptic, and modular structural plasticity during PHP that results in the addition of transsynaptic nanocolumns to the ring.

We did not observe apparent changes in GluR-fluorescence intensity, GluR-cluster number, or homeostatic potentiation of release upon pharmacological GluR perturbation in hypomorphic *neto^109^* mutants. This shows that wild-type Neto levels are required for homeostatic control of GluRs and presynaptic release. GluR inhibition also led to a slight, but significant increase in Brp fluorescence in *neto^109^* mutants, which was less pronounced than in wild type. The defect in PHP seen in *neto^109^* mutants could thus arise from impaired GluR- and/or Brp regulation. Although our genetic data establish a causal relationship between the homeostatic regulation of transsynaptic nanocolumns and presynaptic physiology, future work is required to dissect the mechanisms controlling transsynaptic nanoarchitecture and its homeostatic regulation at this and other synapses.

## Materials and Methods

### Fly Husbandry, Stocks, and Handling

All experiments involving genetically modified organisms have been approved by the responsible authorities (Authorization A120910-4, “Work with GMOs IMLS University Zurich”). *Drosophila melanogaster* strains were reared under standard laboratory conditions and raised at 25°C on standard food. Male and female third instar larvae of the following genotypes were used: *w^1118^*, *BG57-Gal4* (kind gift from Jan Pielage), *UAS-GluRIIA* (*1*), *GluRIIB^SP5^* (this study), and *neto^109^* (*2*) (kindly provided by Mihaela Serpe). *GluRIIB^SP5^* mutants were generated using a CRISPR/Cas9 genome editing strategy as described (*3*). One single guide (sg)RNA line that targeted the sixth exon of the GluRIIB locus (sgRNA: 5’ CATTGATGGATTCTACTCCCGGG 3’) was cloned into the pU6 vector. This construct was sent to BestGene Inc. (Chino Hill, CA) for targeted insertion into the VK18 attP site on the second chromosome. sgRNA flies were crossed to a *vas-Cas9* line on the second chromosome to induce active germline CRISPR mutagenesis, and 20 independent lines were screened by PCR for mutations. This identified 8 independent indel mutations that shifted the open reading frame. *GluRIIB^SP5^* led to an early STOP codon at the 276^th^ amino acid (T276STOP) and was kept for additional analysis.

### Immunostaining

Drosophila larvae were dissected and processed similar as described previously (*4*). In brief, wandering third-instar larvae were dissected in HL3 saline (in mM: 70 NaCl, 5 KCl, 10 MgCl2, 10 NaHCOs, 115 sucrose, 5 trehalose, 5 HEPES, 0.3 CaCl2). After dissection, preparations were washed with HL3 saline and fixed with ethanol (EtOH 100%, Reuss-Chemie, RC-A15-005L) for 10 min. Thereafter, preparations were washed thoroughly with PBS containing 0.1% Triton X-100. For pharmacological glutamate receptor (GluR) blockade (Figure 4 and Figure 5), larvae were either incubated with HL3 (control) or the GluR antagonist philanthotoxin-433 (PhTX; 20 μM) for 5, 15 or 30 minutes at room temperature before applying EtOH. After washing with PBS/Triton X-100, preparations were blocked with 3% normal goat serum in PBS containing 0.1% Triton X-100 for 1 h 30 - 2h. Incubation with primary antibodies was done at 4 °C on a rotating platform overnight. The following primary antibodies and dilutions were used: anti-Bruchpilot (mouse, nc82 (*5*); 1:100), anti-GluRIIC (rabbit, 1:100, kindly provided by Jan Pielage. For stainings in *neto^109^* shown in Figure 5: anti-GluRIIC was used at a dilution of 1:500); anti-GluRIIA (mouse, 1:1000, DSHB), anti-GluRIIB (rabbit, 1:5000, kind gift from Dion Dickman), anti-Neto-β (rabbit, 1:500; kind gift from Dion Dickman). The following secondary antibodies (1:100) were applied for 2 h at room temperature on a rotating platform: Atto 594 (anti-mouse, Sigma-Aldrich, 76085) and Abberior STAR 635 P (anti-rabbit, Abberior, 53399). For stainings in *neto^109^* shown in Figure 5: anti-rabbit STAR 635 P was used at a dilution of 1:250). Preparations were mounted onto slides with ProLong Gold (Life Technologies, P36930). Experimental groups of a given experiment were processed in parallel in the same tube.

### Image Acquisition and Processing

Confocal and gSTED microscopy was performed with an inverse Leica TCS SP8 STED 3X microscope (Leica Microsystems, Germany) of the University of Zurich Center for Microscopy and Image Analysis. For excitation, we used a flexible white light laser with an output range of 470 - 670 nm in combination with a 775 nm STED depletion laser. Excitation light (580nm or 640nm) was focused onto the specimen using a 100x objective (HC PL APO 1.40 NA Oil STED WHITE; Leica Microsystems, Germany) with immersion oil conforming to ISO 8036 with a diffraction index of n=1.5180 (Leica Microsystems, Germany). Emitted light was detected with two HyD detectors in photon counting mode (Leica Microsystems, Germany). For STED imaging, we used time-gated single photon detection (empirical adjustment within a fluorescence lifetime interval from 0.7 to 6.0 ns (*6*)). Pixel size was 10×10nm or 20×20nm and z-stacks were acquired with a step size of 120 or 130 nm. Line accumulation was set to 1 and 6 for confocal and STED imaging, respectively. Images were acquired with LAS X software (Leica Application Suite X, version 2.0; Leica Microsystems, Germany). Experimental groups were imaged side-by-side with identical settings.

Images were processed and deconvolved with Huygens Professional (Huygens compute engine 17.04, Scientific Volume Imaging B.V., The Netherlands). In brief, the “automatic background detection” tool (radius = 0.7 *μm*), and the “auto stabilize” feature were used to correct for background and lateral drift. Images were deconvolved using the Good’s roughness Maximum Likelihood algorithm with default parameter settings (maximum iterations: 10; signal to noise ratio: 7 for STED and 15 for confocal; quality threshold: 0.003). ImageJ (version 1.51n, National Institutes of Health, USA) was used for maximum intensity z-projection.

### Image Analysis

#### Fluorescence intensity and area

Fluorescence intensity and area were analyzed from max. intensity z-projection confocal data using Fiji/ImageJ (version 1.51n, National Institutes of Health, USA). To create a binary mask, max. projection intensity data were background corrected (rolling ball, radius 1 = *μm*) and filtered (3 x 3 median). Thereafter, a threshold was set to 15 - 30% of the maximum fluorescence intensity of a pixel located within the NMJ for the Brp- and GluR channel, respectively. The resulting binary mask was then projected onto the respective background-corrected, unfiltered max. projection data and the intensity and area of individual particles were read out.

#### Ring detection

Ring detection was implemented in C++ using the open source (CeCILL license) “CImg” library (cimg.eu/) for image loading/storage. In brief, the entire deconvolved max. z-projection was scanned pixel-by-pixel in eight directions with regard to a given center pixel with a filter (Figure S2A) to assign each pixel with a ring-center probability (Figure S2B). The filter accumulates fluorescence intensity relative to its center in eight directions (two horizontal, two vertical and four diagonal) within a distance of 40 - 120 nm from the center pixel (Figure S2A). For each pixel, the accumulated values for each direction are added to a score, which is compared to two thresholds. First, the number of directions with filter values larger than the “high threshold” (*th*) is counted. *th* was defined as the maximum score multiplied by an empirically derived constant of 0.05 (see also below, “*Line profiles*’”). Second, the number of directions with filter values larger than the low threshold (*tl*, defined as the maximum score multiplied by a constant of 0.025) is counted. Based on the eight directions, both values are within a range from zero to eight. A ring-center probability larger than zero is assigned to a pixel if both values are larger than six. The threshold of six is based on calibration experiments in which we systematically changed the threshold and visually inspected the results. Pixels with the highest ring-center probability are first assigned as “ring centers”. If two pixels with ring-center probabilities greater than zero are closer than 10 pixels (200 nm) distance, only the pixel with the higher probability is assigned as a ring center. Note that the algorithm considers local fluorescence gradients rather than a threshold.

#### Line profiles

Line profiles with a length of 600 nm were either independently anchored in Brp- and GluR-ring centers (Figure S2D), or in Brp-ring centers alone (Figure S2D; Figure 1 – 5). For each line profile, we read out the Brp- and GluR fluorescence intensity along a line that was rotated by 360 degrees in single degree steps, and centered at the respective ring center. Normalized line profiles were obtained by dividing the profile of each ring by its fluorescence intensity maximum. Based on different fluorescence maxima locations within the ring, the peaks of the average line profiles are smaller than one. Average line profiles are based on all rings detected in a given experimental group. Line profiles with a fluorescence maximum within a diameter of <100nm with respect to the ring center were defined as “not ring-like” line profiles (Figure S2F). The ring detection algorithm was adjusted such that the fraction of “not ring-like” line profiles was close to zero in the Brp channel. For the GluR channel, the fraction of “ring-like” line profiles for the respective genotypes is given in Figure S2F.

#### Local maximum detection

For the detection of local maxima, i.e. ‘nano-clusters’ within a ring (Figure 1D’, 1E, Figure S3A, S3B), two criteria were applied: The first criterion assigns high cluster-center probabilities to pixels with high fluorescence intensity relative to neighboring pixels within a distance of one to three pixels (Figure S3A). The second criterion is in place to evaluate adjacent pixels and to exclude ‘isolated’ pixels with high fluorescence intensity. Pixels with fluorescence intensities above a threshold of 20 a.u. (8-bit) get initialized with the maximum cluster-center probability, while pixels with intensities below this threshold get assigned with a cluster-center probability of zero. Then, the fluorescence intensity ratios between nearby pixels around a potential cluster center are analyzed. Nearby pixels in a range from one to three are considered according to the following scheme (Figure S3A): Depending on the distance to the center, these nearby pixels are assigned with different intervals (distance of one pixel: 0.60 - 0.97; distance of two pixels: 0.32 - 0.98; distance of three pixels: 0.01 - 0.99). If the calculated intensity ratio of a nearby pixel to the center pixel is within the specific interval, the cluster-center probability of the center pixel is not decreased. If the calculated intensity ratio is outside the interval, the probability of the center pixel is decreased by the distance of the ratio to the nearest point of the interval and multiplied by a factor 50. For all pixels with a cluster-center likelihood greater than a threshold of 20 a.u., a second criterion is applied in two steps as follows: First, if any pixel within a distance of one or two in x or y direction shows a higher intensity than the center pixel, the respective center pixel is no longer considered as a potential cluster center. If e.g. two pixels next to each other with a symmetrical intensity decrease in the nearby region have a cluster center probability greater than zero assigned from the first criteria, only the brighter pixel keeps its cluster center likelihood, while the other pixel gets assigned with a likelihood of zero. Second, if more than three of the eight pixels surrounding the center pixel have an intensity below 10 a.u., the center pixel gets assigned with a cluster center probability of zero. With this rule, isolated bright pixels surrounded by very low-intensity pixels are excluded as cluster centers. On our images, these cases most likely represent noise, as they are evenly distributed over the whole image. Finally, pixels with the highest cluster-center probability get assigned as cluster centers first. Subsequently, pixels with lower cluster-center probabilities get assigned as cluster centers. If two pixels with cluster-center probabilities greater than zero are closer than 3 pixels distance, only the pixel with the higher probability gets assigned as a cluster center. This fluorescence intensity gradientbased approach allows detecting local maxima in heterogeneous fluorescence intensity landscapes (Figure S2A).

### Electrophysiology

Wandering third-instar larvae were dissected in HL3 solution (5 mM KCl, 70 mM NaCl, 10 mM Na-HEPES, 5 mM HEPES, 5 mM trehalose, 115 mM sucrose, 10 mM MgCl2) with 0.35 or 0.8 mM CaCl2 for sharpelectrode membrane voltage recordings. The internal organs, including brain and ventral nerve cord, were carefully removed from the body-wall with intact muscle fibers and innervating motor nerves. Sharp-electrode recordings were performed on muscle 6 of segments 3 and 4 with sharp glass electrodes (resistance 10-25 MΩ) using an Axoclamp 900A amplifier (Molecular Devices). For individual NMJs, mEPSPs were recorded prior to EPSPs induced by stimulating the respective hemi-segmental nerve with single APs (3 ms stimulus duration for EPSPs, 0.3 Hz). A total of 50 EPSCs were recorded to obtain the mean EPSP amplitude for each cell.

Semi-intact (dorsally dissected; with internal organs, brain and ventral nerve cord intact) larvae were incubated with GluR antagonist PhTX-433 (20 μM; Cat # sc-255421, Santa Cruz Biotechnology) for ~15 minutes. This was followed by HL3 washes, removal of internal organs, brain and ventral nerve cord to obtain a fully-stretched preparation for electrophysiological recordings.

### Quantification and Statistical Analysis

Data were analyzed using Prism (GraphPad Software, CA, USA) and IgorPro (WaveMetrics Inc, Lake Oswego, OR, USA; v6.37) and tested for distribution via D’Agostino-Pearson omnibus normality test. Line profile intensities are represented as mean ± SEM and line profile intensities in Figure 1C, Figure 2 (D, H, I, M, N), Figure 3 (D, H, I), Figure 4 (H, I), Fig 5. (L, M) and Figure S5 were analyzed through Kruskal-Wallis test with Dunn’s Post Hoc multiple comparison test. Cluster counts in Figure 2 (E, J, O), Figure 3 (E, J), Figure 4J and Figure 5N are shown as box-and-whisker plots with median ± min to max and, unless otherwise stated, they were analyzed through Kruskal-Wallis test with Dunn’s Post Hoc multiple comparison test. Exceptions: Unpaired Student’s t-test with Welch’s correction (1E and 2E), Mann-Whitney test (3E), Ordinary one-way ANOVA with Bonferroni’s Post Hoc multiple comparison test (3J), one sample t test (hypothetical value: 100; Figure S6). Cumulative frequency distributions were analysed via Kolmogorov-Smirnov test (4C, 4D, 5G, 5H). PhTX-induced Brp intensity changes in *neto^109^* compared to *w^1118^* (reported in the main text) were tested through Mann-Whitney test. Sample size and P-values are reported in the text and in the supplementary tables. Unless otherwise noted, *n* refers to the number of synapses as identified by Brp rings. Unless otherwise noted, data are based on *N* ≥ 4 NMJs. Reported sample size and corresponding statistic tests were based on “n” defined as “active zone (AZ)/ring” number. We ran a linear mixed model (Multicomp package in R; version 3.5.3) to test if the statistical differences of the recorded parameters depend on our sample size definition by considering NMJ number (N) and AZ number (n) as random effects, and treatment condition (HL3 vs. PhTX) as the fixed effect for our largest data set (ring diameter and cluster number, 30’ HL3 vs. PhTX, Figure 4). The results of the linear mixed model suggest that the statistical differences observed in this data set are due to the fixed effect (treatment) rather than the random effect (N vs. n), suggesting that our conclusions are independent of our sample size definition.

## Acknowledgments

We are grateful to Igor Delvendahl, Anu Nair, Olivier Urwyler and Dietmar Schmucker for critical comments on previous versions of this manuscript. We would like to thank Jan Pielage for sharing anti-GluRIIC antibody, and Mihaela Serpe for providing the *neto^109^* mutants. Many thanks to the members of the Center for Microscopy and Image Analysis of the University of Zurich, especially to Jana Döhner, for technical support.

## Funding

This study was supported by an SNSF-professorship grant (PP00P3_144816) and an ERC Starting Grant (‘SynDegrade’, 679881) to M.M.

## Author contributions

P.F., P.M. and M.M. conceptualized and designed the study, P.F. wrote analysis code, P.F. and P.M. acquired data, P.F., P.M. and M.M. analyzed data, P.M. and M.M. wrote the manuscript.

## Competing interests

The authors declare no competing interests.

## Data and materials availability

All data, code, and materials underlying the findings of this study are available from the corresponding author upon reasonable request.

## Notes

### Competing Interest Statement

The authors have declared no competing interest.

